# Visibility graphs for fMRI data: multiplex temporal graphs and their modulations across resting state networks

**DOI:** 10.1101/106443

**Authors:** Speranza Sannino, Sebastiano Stramaglia, Lucas Lacasa, Daniele Marinazzo

## Abstract

Visibility algorithms are a family of methods that map time series into graphs, such that the tools of graph theory and network science can be used for the characterization of time series. This approach has proved a convenient tool and visibility graphs have found applications across several disciplines. Recently, an approach has been proposed to extend this framework to multivariate time series, allowing a novel way to describe collective dynamics. Here we test their application to fMRI time series, following two main motivations, namely that (i) this approach allows to simultaneously capture and process relevant aspects of both local and global dynamics in an easy and intuitive way, and (ii) this provides a suggestive bridge between time series and network theory which nicely fits the consolidating field of network neuroscience. Our application to a large open dataset reveals differences in the similarities of temporal networks (and thus in correlated dynamics) across resting state networks, and gives indications that some differences in brain activity connected to psychiatric disorders could be picked up by this approach.

## Introduction

Visibility graphs (VG) were recently introduced as a method to map time series into networks [29, 39], with the aims of using the tools of Network Science [5, 43] to describe the structure of time series and their underlying dynamics. This strategy of transforming time series into graphs has been exploited in recent years by some authors and several alternative methods have been put forward, contributing to the nascent field of performing graph-theoretical time series analysis (see [12, 62, 64] for a few seminal examples and [15] and references therein for a recent overview). Research on VG has since then focused essentially on two separated avenues. First, analytic studies have primarily explored the foundations of this mapping [14, 22, 27, 38] and elaborated on mathematical methods [26] to extract rigorous results on the topology of visibility graphs associated to canonical dynamics such as stochastic or chaotic processes [6, 16, 30, 37, 40] and to obtain combinatoric analogues of different dynamical quantities [32]. The second avenue deals with applications of this machinery, primarily by using this method as a feature extraction procedure with which build feature vectors which can properly characterize time series with the purpose of making statistical learning (see [4, 20, 36, 53] for a few examples in the life sciences). In this latter context, the application to neuroscience is on its infancy, and has essentially limited so far to the analysis of electroencephalogram (EEG) data (see [1–4, 42] for a few examples). The study of fMRI recordings under these lens has been scarce, and in this work we would like to motivate and justify why we think this is a promising enterprise, both from a univariate and -perhaps more interestingly-from a multivariate time series perspective [31]. Among other strategies to map time series intro graphs, using the repertoire of visibility graphs is particularly interesting, not just because its current application is scarce, but also because these methods are well suited to handle the specificities of fMRI data. More concretely, these methods have been shown to be efficient in extracting information and dealing with (i) data polluted with noise [30], (ii) multivariate [44], and (iii) non-stationary [39]. In order to showcase the usefulness of visibility graphs in neuroscience we will choose a biggish, high quality public dataset of resting state fMRI data [45], and will make use of the family of visibility algorithms to build a multilevel graph of temporal networks, where each node represents a time point, and two nodes are connected if they are *visible* to each other, according to the algorithm explained below. In the case of multivariate time series -as the ones acquired in neuroimaging-each of these networks is actually the layer of a *multiplex* network (usually associated to a recording in a different ROI). By being able to integrate in a single structure all the data enables both the intralayer (univariate) and the interlayer (multivariate) analysis simultaneously. We will show that a direct analysis of this network provides genuine and nontrivial information on fMRI data, potentially including -but not only- the description and possible non-invasive classification of some brain diseases.

## Materials and Methods

### fMRI data

We used the public dataset described in [45]. This data was obtained from the OpenfMRI database, its accession number being ds000030. We use resting state fMRI data from 121 healthy controls, 50 individuals diagnosed with schizophrenia, 49 individuals diagnosed with bipolar disorder and 40 individuals diagnosed with ADHD. The demographics are reported in the original paper, and they can additionally be found in the GitHub page containing the results of this study^1^.

The fMRI data was preprocessed with FSL (FMRIB Software Library v5.0). The volumes were corrected for motion, after which slice timing correction was applied to correct for temporal alignment. All voxels were spatially smoothed with a 6mm FWHM isotropic Gaussian kernel and after intensity normalization, a band pass filter was applied between 0.01 and 0.08 Hz. In addition, linear and quadratic trends were removed. We next regressed out the motion time courses, the average CSF signal and the average white matter signal. Global signal regression was not performed. Data were transformed to the MNI152 template, such that a given voxel had a volume of 3mm x 3 mm x 3mm. Finally we averaged the signal in 278 regions of interest (ROIs) using the template described in [54].

In order to localize the results within the intrinsic connectivity network of the resting brain, we assigned each of these ROIs to one of the 9 resting state networks (7 cortical networks, plus subcortical regions and cerebellum) as described in [57].

### Construction of the visibility graphs

The procedure to build up a visibility graph is extensively and clearly described in [29, 30, 32] for univariate and [31] for multivariate time series. Here we will recall the basic steps and provide a visualization of the application of the methodology to BOLD data.

Given a time series of *N* data, any two time points *i* and *j* in which the measured quantity takes the values y and *y_j_* respectively will have visibility and consequently will become two connected nodes in the associated *natural visibility* graph if any other data point *y_k_* placed between them fulfills the condition:

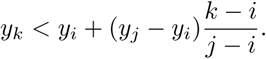

Together with this convexity criterion, named *Natural Visibility* (NV), an ordering criterion, named *Horizontal Visibility* (HV) has also been defined [30]. According to the latter, two time points *i* and *j*, in which the measured quantity takes the values y*_i_* and *y_j_* respectively, will now have horizontal visibility if any other data point *y_k_* placed between them is smaller, i.e.
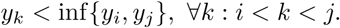

In either case, the resulting graphs have *N* nodes, are connected by a trivial Hamiltonian path that induces a natural ordering in the degree sequence, and are undirected (see figure 1 for an illustration). In the event the the time arrow turns out to be a relevant aspect, directed graphs can be easily constructed, as detailed in [32]. Note that the resulting Horizontal visibility graph (HVG) is simply a core subgraph of the Natural visibility graphs (NVG), the former being analytically tractable [26]. As a matter of fact, HVG can be understood as an order statistic [28] and therefore filters out any dependency on the series marginal distributions (that’s not true for NVG so in applications where marginal distributions are relevant, one should use NVG over HVG).

**Figure 1.**
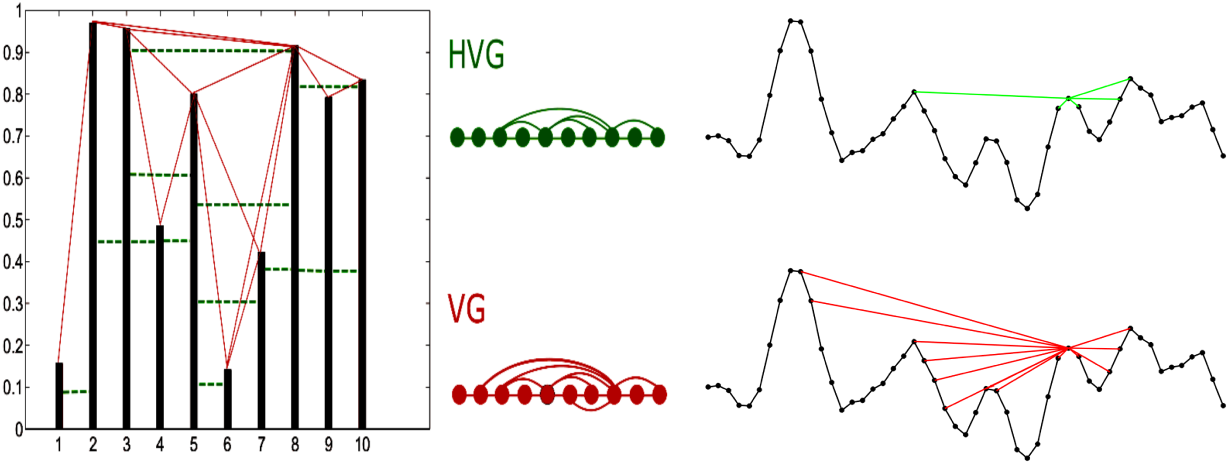
Examples of Natural Visibility Graph (VG, bottom) and Horizontal Visibility Graph (HVG, top) algorithms applied to the same sample time series. In each case, a time series of *N* data map into a graph of *N* nodes, where two nodes are linked according to either *natural* or *horizontal* visibility criteria (i.e. convexity and ordering criteria respectively, see the text). On the right side, an illustration of the points connected according to either criterion to a given time point from a typical fMRI region of interest time series.

Both algorithms are fast: naive implementations of NVGs have a runtime complexity O(N^2^), however a divide-and-conquer strategy already reduces it to O(*N* log*N*) [33]. Naive implementation of HVG is already O(*N* log*N*) in most of the cases of practical interest. Finally, these methods are well-suited to handle several degrees of non-stationarity in the associated time series [28].

In this work we will be analyzing BOLD data, and for that task we decided to choose NVG over HVG. This is because NVGs are in principle better suited to handle and extract long range correlations than HVG, as the former naturally allow for the development of hubs -which will be typically associated to extreme events in the data and can correlate with data at all scales-. Correlations in time series are actually inherited in graph space in the degree distribution. It is somewhat easier to find fat-tailed degree distributions in NVGs (which account for hubs with extremely large degrees). On the other hand, HVGs (which have shown to work fine with processes evidencing short-range correlations) typically display exponentially decaying degree distributions: a feature which is linked to short-scale visibility, making this method more local.

For illustration, Figure 1 depicts how the links are established in the visibility graph according to both visibility criteria. The code used to compute the Visibility Graphs is available ^2^, and is basically a translation to Matlab of the original visibility scripts in Fortran90 ^3^.

When it comes to the application to multivariate time series formed by M series, note that each of the M time series yields a different visibility graph to begin with, so in principle the multivariate series can always be mapped into a multilayer graph with M layers [31]. Moreover, since for every node *i* there is a natural correspondence across layers (node *i* corresponds to time stamp *i* and this is the same time stamp for all components), there exist a natural alignment between every node of each layer, so the multilayer graph is effectively a so-called multiplex network [5, 31] (see figure 2 for an illustration). Of course, other smarter alignments between graphs could be investigated (for instance, one could try to find the alignment that minimize some sort of Hamming distance between ordered node sets), but in this work we keep it simple and consider the natural alignment induced by the time arrow.

Interestingly, this multiplex visibility graph encodes the complex structure of each time series in the topology of each layer. One can therefore extract in each layer any desired topological feature (say for instance the entropy over the degree distribution, which would provide a different number for each layer), with which one could build a feature vector that provides a compact representation of the multivariate time series complexity. A similar procedure was followed for instance in [1] to extract markers of Alzheimer’s disease from a graph theoretical characterization of the Hurst index of EEG data.

**Figure 2.**
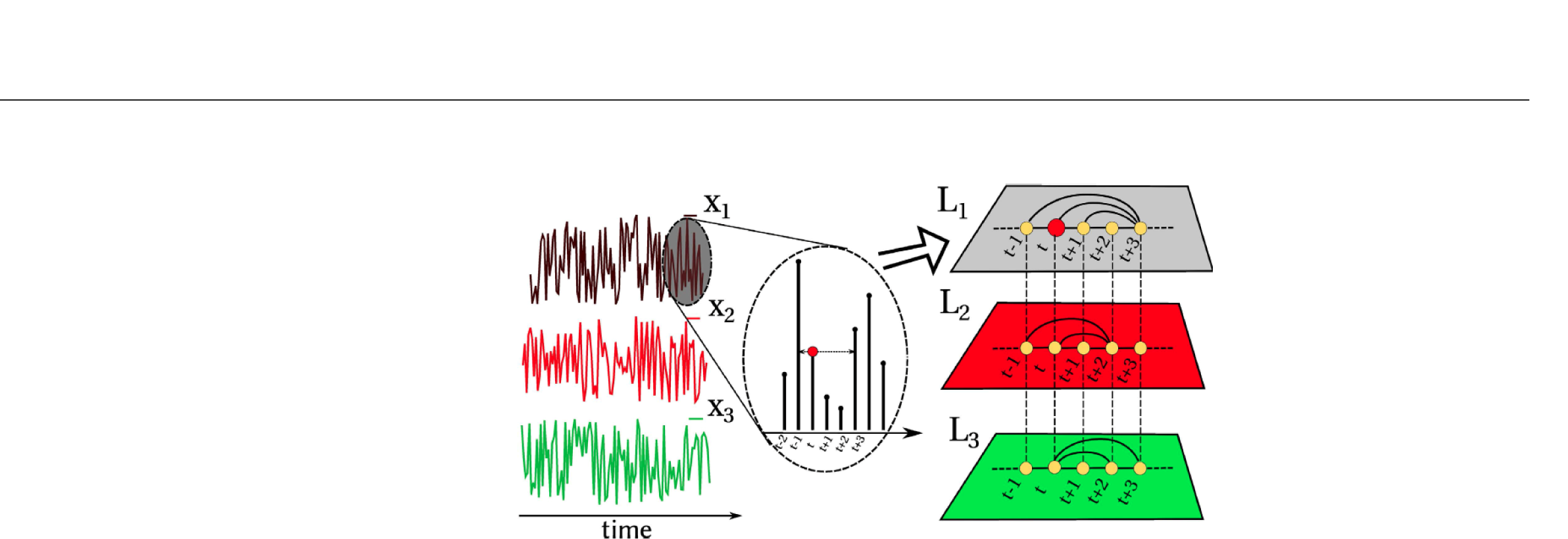
Example of the construction of a multiplex visibility graph from a multivariate time series with *M* = 3 components. In this cartoon, each layer builds the HVG associated to each variable, therefore all layers are well aligned according to the time arrow, making interlayer comparison straightforward. Adapted from [31].

Second, the complex interdependencies and correlations which might emerge in a multivariate series across variables could in turn be extracted using similarity measures across layers. There exist a large variety of network measures that one can use for this task [44]. A simple example of such a measure is the so-called interlayer mutual information, recently explored in the context of multiplex visibility graphs of coupled chaotic maps [31]. This quantity measures the information shared by every two layers based on the similarity of the degree distributions. Given the degree distributions *P*(*k^α^*) and *P*(*k^β^*) of two arbitrary layers *α* and *β* it is defined as

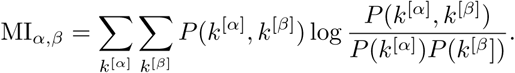

As the degree distribution captures the structure of each layer, this measure is in turn capturing the information shared between the two layers, that is to say, the information shared across each time series component of the multivariate time series. Now, since this is a *M* × *M* matrix whose *ij* entry provides the mutual information between layers (ROIs) *i* and *j*, one can then -for instance- average across pairs (that is to say, across ROIs) to find a scalar quantity 〈MI〉: the mean value of the mutual information for each intrinsic connectivity network. This methodology is depicted in figure 3. Note that other informational or similarity measures between layers could be used instead (e.g. edge overlap, conditional or partial mutual information, transfer entropy, etc), here for the sake of exposition we only consider mutual information.

The visibility algorithms produce networks whose nodes are time points. As one can observe in figure 3, these networks have a modular structure, in which subnetworks are constituted by time points that are mainly adjacent. A modular structure in a temporal network is thus an indication of different temporal regimes. The existence of these temporal regimes is what motivated the study of dynamical functional connectivity (see for example [17, 21]). Dynamic functional connectivity can be seen in the visibility framework as the comparison of the temporal networks, taking their modular structure into account. This comparison can be done in the first place considering the modular network as a whole. In our case we partitioned the visibility graphs for each ROI using hundred runs of the Louvain algorithm. We then quantified the distance between the two partitions by means of the mutual information, using the function in the Brain Connectivity Toolbox [50]. The results of the partition of two ROIs, one in the Anterior Cingulate Cortex (ACC) and one in the Precuneus (PCC), are shown in figure 4. The modules of the graphs correspond to consecutive time points (left panels), i.e. partitioning the visibility graph provides a natural decomposition of the time series in time intervals. Turning to the interdependency between the two time series, the right panel of figure 4 represents the Sorensen similarity between each pair of modules in the two time series. It shows that there are segments with high Sorensen indexes, and it is likely that during these segments the two ROIs reflect similar neural events.

**Figure 3.**
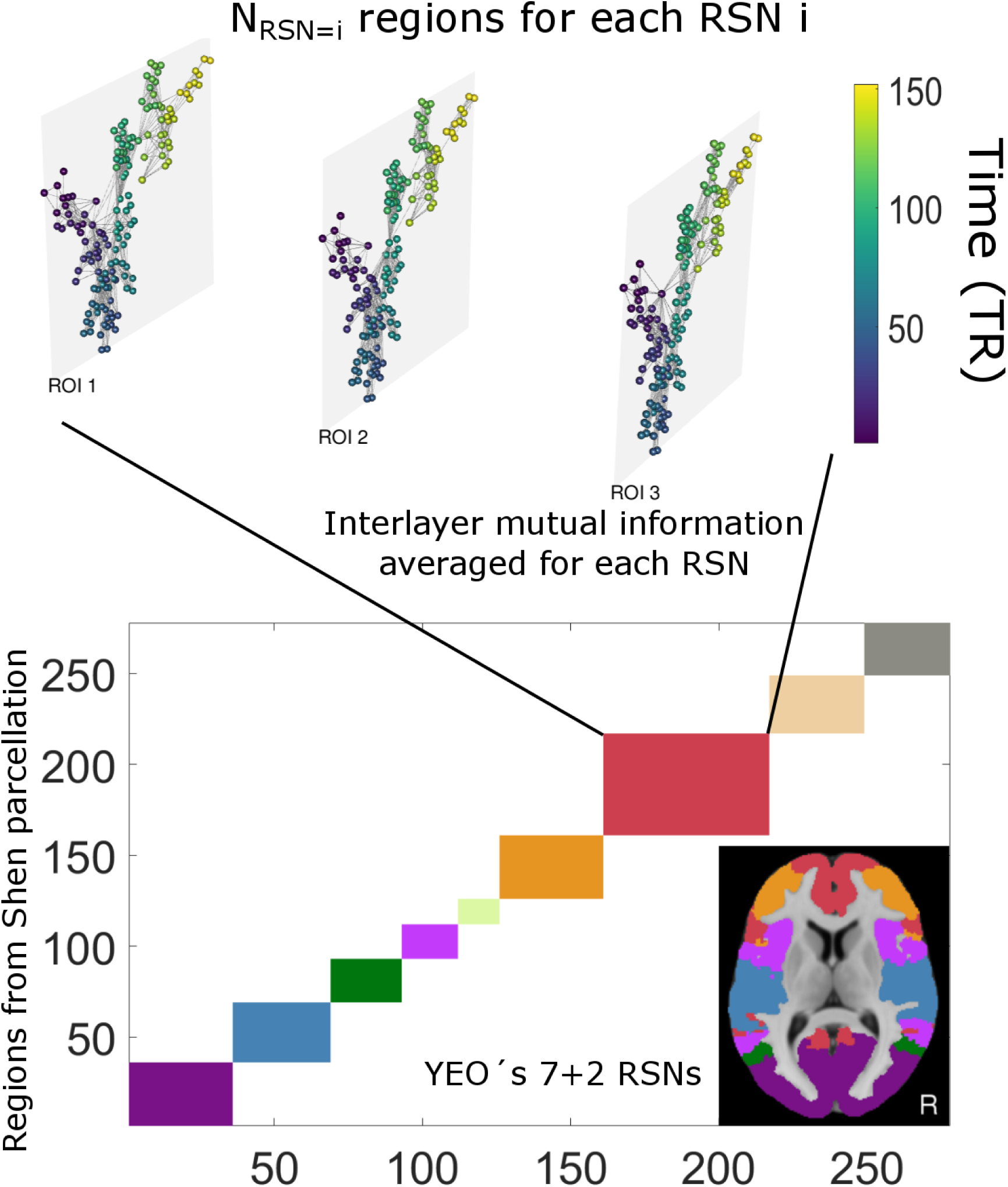
Scheme of the procedure: Within a given region which aggregates a certain number of ROIs, one constructs a visibility graph per ROI and builds accordingly a multiplex visibility graph. We then compute the pairwise mutual information between degree distributions across the multiplex layers (ROIs) and finally average to obtain a value for each RSN. The multilayer network is visualized with MuxViz [10]

**Figure 4.**
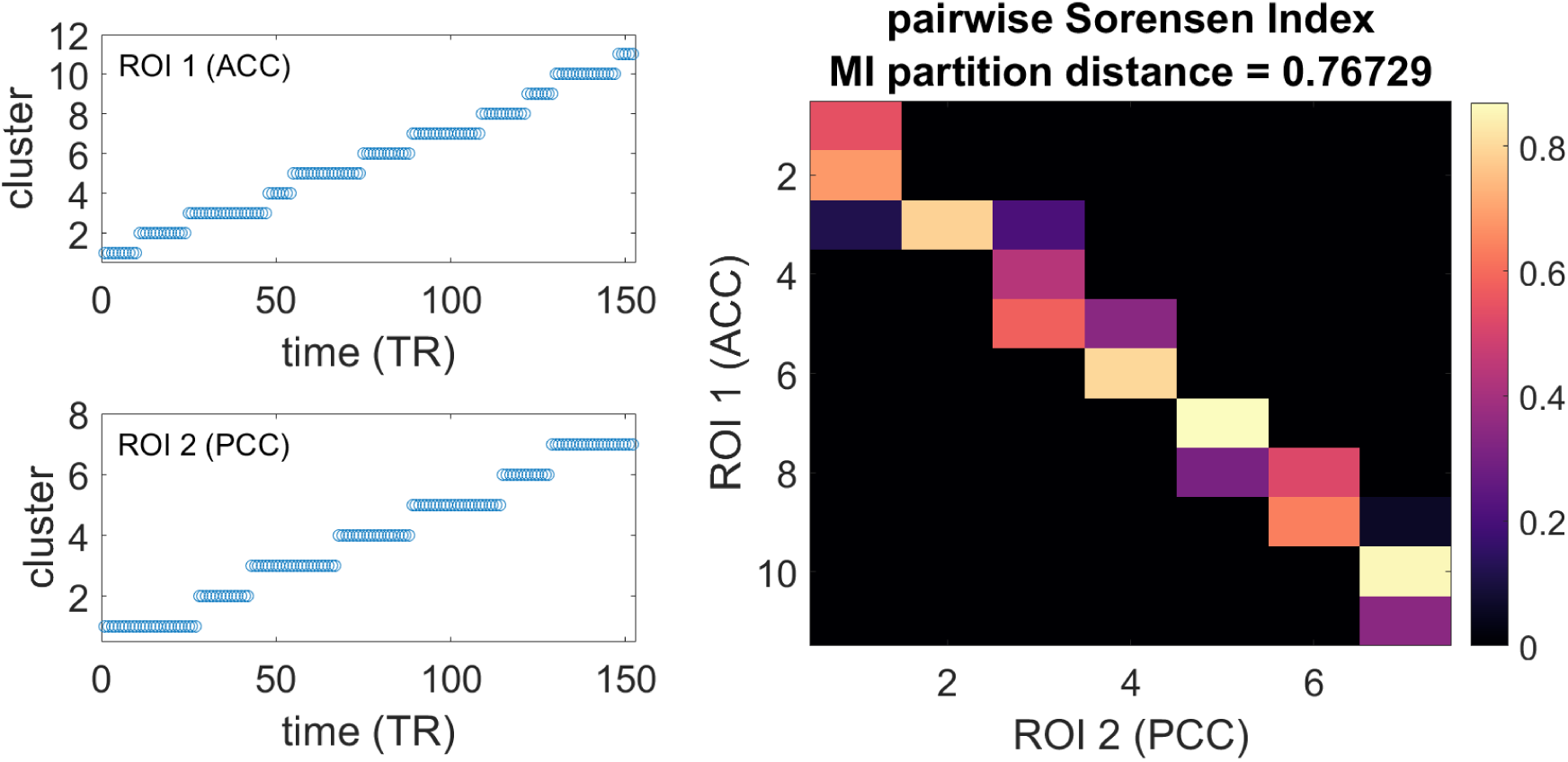
Left: The clusters in which the visibility adjacency matrices from two example ROIs are partitioned according to the Louvain algorithm. Right: Sorensen index quantifying the similarity between pairs of clusters. The value of the distance among the partitioned networks considered as a whole is also reported, in terms of normalized mutual information.

## Results

We start by reporting in Figure 5 the results of 〈MI〉 within each of the intrinsic connectivity networks, for the four groups of subjects considered. For each group of subjects, each circle corresponds to 〈MI〉 of a given subject, and Random Average Shifted Histograms (RASH) are also provided. This representation is not parametric, and it is bounded. The plots report the median of the Harrell Davis estimator, and the 95% high density intervals using a Bayesian bootstrap. The outliers are detected based on the distance between each pair of data points without assuming symmetry of distributions.

In order to account for departure from normality of these distributions we used a graphical approach and computed the Kolmogorov-Smirnov distance, using the publicly available code ^4^, obtaining values up to 0.7 (a value of 0.39 would correspond to rejecting the null hypothesis at a level *α* < 0.001 for the smallest population). The number of ROIs constituting each intrinsic state network (thus a proxy for the network size, given that Shen’s parcellation has ROIs of similar size) is not correlated with the average value of the mutual information. In particular, it is interesting to observe that the intrinsic connectivity network called *Limbic* in Yeo’s parcellation is the smallest one, but nonetheless has a low interlayer mutual information compared to the other networks for all the clinical groups.

The network which showed a clearest differentiation in terms of the average interlayer mutual information among the four clinical groups is indeed the *Limbic* one (Figure 6). This evidence was assessed by means of a multivariate response test with age of the subjects and framewise displacement as covariates. The p-value of 0.005 was corrected for multiple comparisons using the Bonferroni-Holm criterion with *α* = 0.05. The Kolmogorov-Smirnov statistics of the pairwise comparison between the distributions of average interlayer mutual information values for these particular networks ranged from 0.15 to 0.3. The null hypothesis of values for controls and schizophrenics drawn from the same distribution would be rejected with an *α* < 0. 005. Figure 6 also reports the shift functions to visualize the difference between two distributions, in this case controls and schizophrenics. This function [59] does not assume (as t-tests do) that two distributions differ only in the location of the bulk of the observations, and allow to determine how, and by how much, two distributions differ. Here the Harrell-Davis quantile estimator is used. Confidence intervals of the decile differences with a bootstrap estimation of the standard error of the deciles are computed, and one controls for multiple comparisons so that the type I error rate remains around 0.05 across the 9 confidence intervals ^5^. In this specific case we can observe a clear separation for all the quantiles but the ninth one.

**Figure 5.**
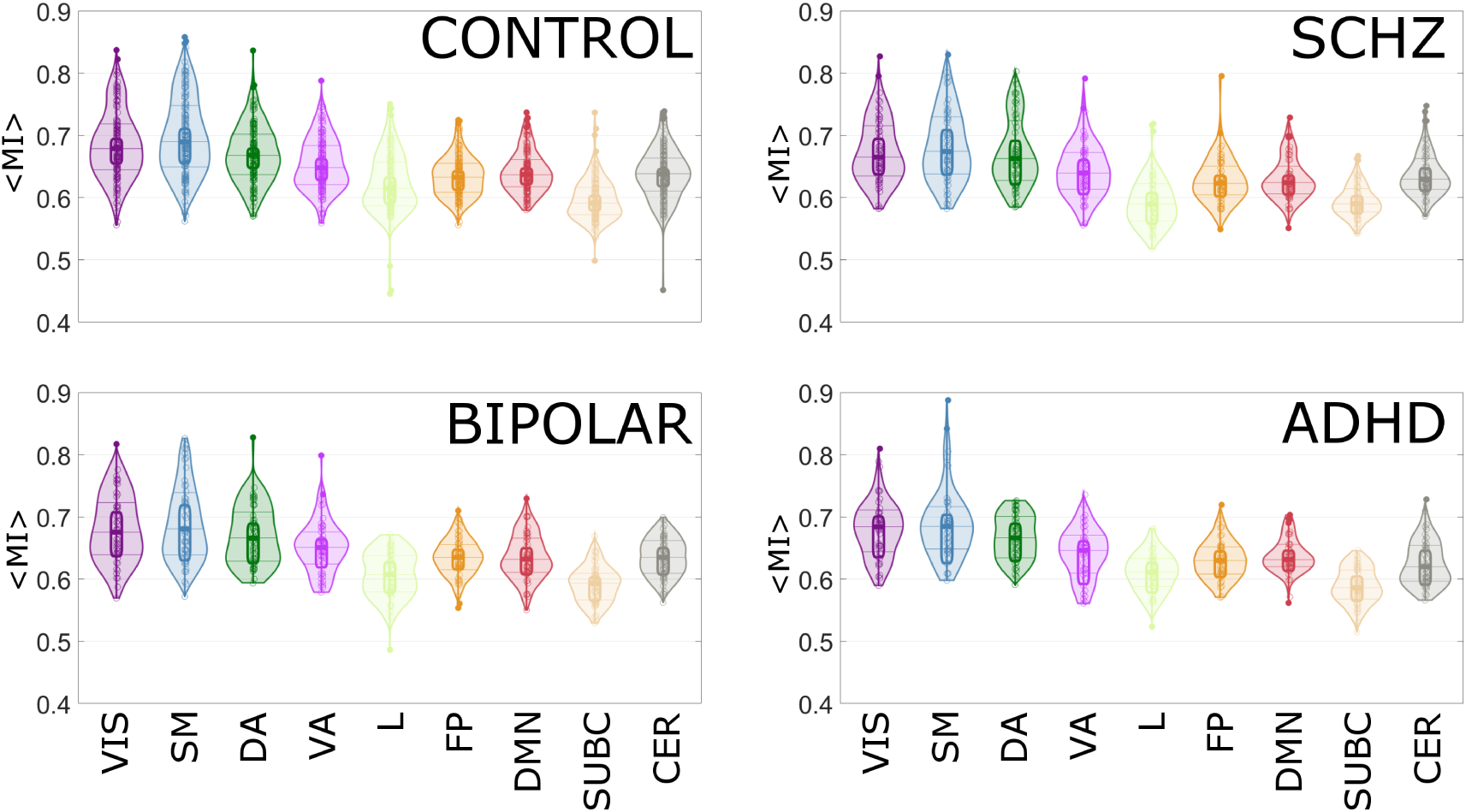
For each group and each intrinsic connectivity network, we plot the distribution across subjects of the averaged interlayer mutual information.

To complement this analysis, in figure S1 we further report two additional ways in which results of this kind are often represented (mean and standard errors). According to this plot it is already evident to a naked eye that the method easily distinguishes controls from patients with any mental disorder, suggesting that visibility graphs do indeed extract informative features which can be used for non-invasive diagnosis. It shall be stated that visualizing results in such a way is indeed suboptimal and sometimes problematic (nicely explained in [49]), that’s why we initially chose the visualizations provided in figures 5 and 6.^6^,^7^

**Figure 6.**
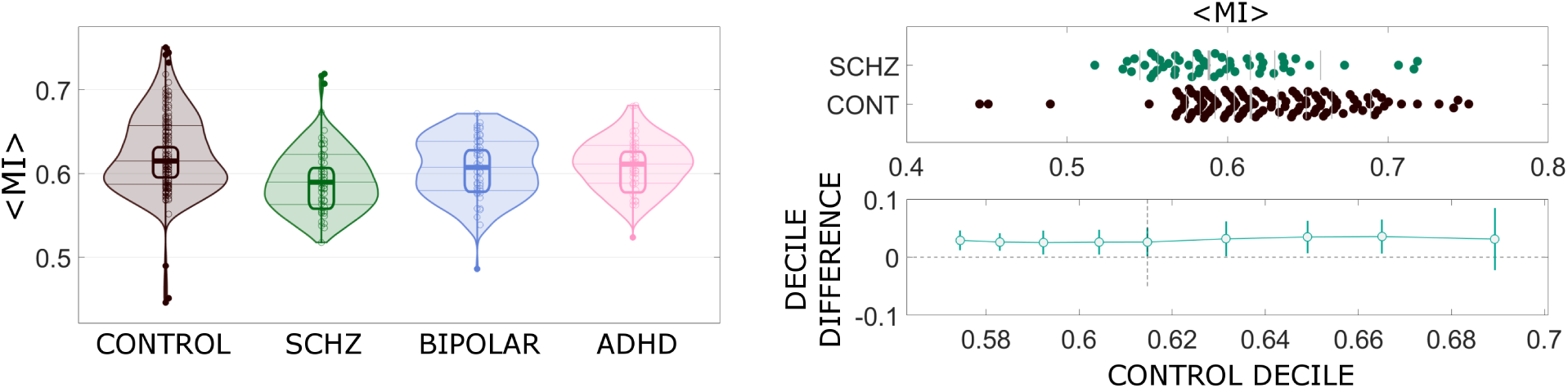
Left: The average interlayer mutual information for the intrinsic connectivity network denoted as *Limbic*, for the four groups of subjects. Right: Shift function to visually and statistically compare the distributions for controls and schizophrenics, at different quantiles

## Discussion

### Why the (Multivariate) Visibility Graph?

All in all, there are several reasons why we think that visibility graphs are a convenient tool, here we discuss some.

#### Usefulness

Visibility Graphs have been shown to inherit in their topology the essence of the associated dynamics, including nontrivial fingerprints which turn to be both descriptive and informative for statistical learning purposes.

#### Fit for purpose

These methods can be used directly in both stationary and non-stationary signals (i.e. non-stationarity is not required to be removed). Also, series do not require *ad hoc* phase partitioning or symbolization. Also, visibility graphs naturally filter out linear trends so they don’t require such detrending [29]. Furthermore, since HVG is an order statistic, it is also invariant under monotonic (order-preserving) rescaling on the data [28]. The NVG is not invariant under this latter transformation though, so nonlinear rescaling to make data more ‘peaky’ will necessarily modify the associated NVG in a nontrivial way.

#### Computationally easy and efficient

The method is numerically straightforward to implement and the runtime algorithms are quite decent (varying from *O*(*n*) for so-called visibility sequential motifs [22] to *O*(*n* log *n*) for the full adjacency matrices using divide-and-conquer strategy).

#### Amenable to analytical insight

Differently from other strategies for graph-theoretical time series analysis, Visibility Graphs are not computational black boxes. More particularly for the HVG (but not only [22, 39]), there exist several theorems available and methods to build rigorous results of HVGs properties [26, 27, 30, 32, 37], this latter being an are of intense research activity at the interface between combinatorics and dynamical systems. *Versatile:* The methods are not context dependent but are generally applicable to both univariate and multivariate time series across the disciplines. A drawback of this property is that the topological features one can extract from these graphs are themselves not context-dependent.

#### Novel

It builds a bridge between time series and networks and thus opens the exciting possibility of exploring the usefulness of a large bunch of new tools in the endeavor of describing and classifying complex signals.

Coming back to the specific reason why we think that Natural Visibility Graphs are particularly suited for BOLD data, it has been shown that relevant information on the time course of the BOLD signal and on correlated activity can be extracted by looking at single frames, corresponding to peaks in the signal [35, 55], and that these events could be the proxy for an innovation signal at the neural level [23, 60]. In this framework, the degree of the nodes corresponding to the BOLD peaks in the adjacency matrix constructed according to the Natural Visibility emphasizes the functional relevance of the neural events and of the corresponding patterns of coactivation across the brain. Notwithstanding, both NVG and HVG have been shown to be useful in different contexts so there is no general rule of thumb on what method should we use: this choice shall be addressed in a case by case basis. Finally, what is important/informative when it comes to describe the properties of a certain cognitive state? Is it the complex pattern underlying the structure of individual time series (that is, local activity of ROIs) of different regions? Or are the correlations and interdependencies (understood in a broad sense) between these regions the key aspect to look at? When the latter is the case, then a functional network analysis approach [7] seems to be the appropriate thing to do. In the former case where the nature of local activity across regions already captures information [18, 63], then one does not need to resort to functional dependencies and local analysis is the correct thing to do. This is obviously an open question which should be addressed, from a biological point of view, in a case by case basis. A recent study suggests that both conceptual frameworks can indeed be connected [52]. In general, most probably both aspects play a relevant role, and some studies have already successfully merged the two [9, 56]. Be that as it may, the multiplex visibility framework offers a compact way of extracting at once both the local temporal structure (via the network intralayer properties) and the global interconnection pattern (via multiplex interlayer similarities).

### Similarity with other measures

We have discussed at the end of the Methods section that the modular temporal graphs resulting from the visibility algorithm are a natural way to describe different dynamical regimes of individual time series, and their interdependence, without arbitrary and possibly problematic choices such as a sliding window and its length [19, 25].

Features of the visibility graph, such as the modularity, the clustering coefficient, or the node degree could be used as features in classification algorithms aimed to detect modulations of the local and correlated dynamical regime of BOLD signals.

Furthermore, using the excellent resource that is NeuroVault (http://neurovault.org/) we also looked at the maps depicting the results of other measures, and noticed that the areas belonging to the *limbic* Yeo network are associated with lower levels of Regional Homogeneity (ReHo) [63], higher coefficient of variation of the BOLD signal [61], and lower value of the fractional amplitude of low-frequency fluctuations (fALFF) [65]. This evidence speaks to the fact that interlayer mutual information in multiplex visibility networks is associated to decreased predictability and increased independence between the degrees of freedom of the measured time series.

### Classification of neural disorders

The main focus of this paper is methodological, and a thorough discussion on the implications of our results on neuroimaging studies of psychiatric disorders is beyond its scope, moreover we wouldn’t want to hypothesize after the results are known (HARKing [46]). That being said, it is interesting to highlight that the limbic network has been previously associated to mental disorders [24, 34, 47, 48, 51, 58]. In the same way we refer the reader to nice recent studies specifically aimed to use advanced neuroimaging data analysis tools to map and classify neural disorders [8, 11, 41], and [13] for a review. Our results shown here using visibility graphs confirm some of these previous works and further showcase that (i) visibility graphs extract informative features with which (ii) we can find statistically significant signatures of different neural disorders.

To conclude, given the exposition and results reported in this study, we hope to have motivated our colleagues to consider Visibility Graphs as a valuable Network Neuroscience tool for both exploratory and focused studies.

## Supporting Information

The code and data necessary to replicate the results reported here are indicated in the text. For convenience we report here the location of the main repository, linking to the others.

https://github.com/danielemarinazzo/Visibility_LA5C_data

**Figure S1.**
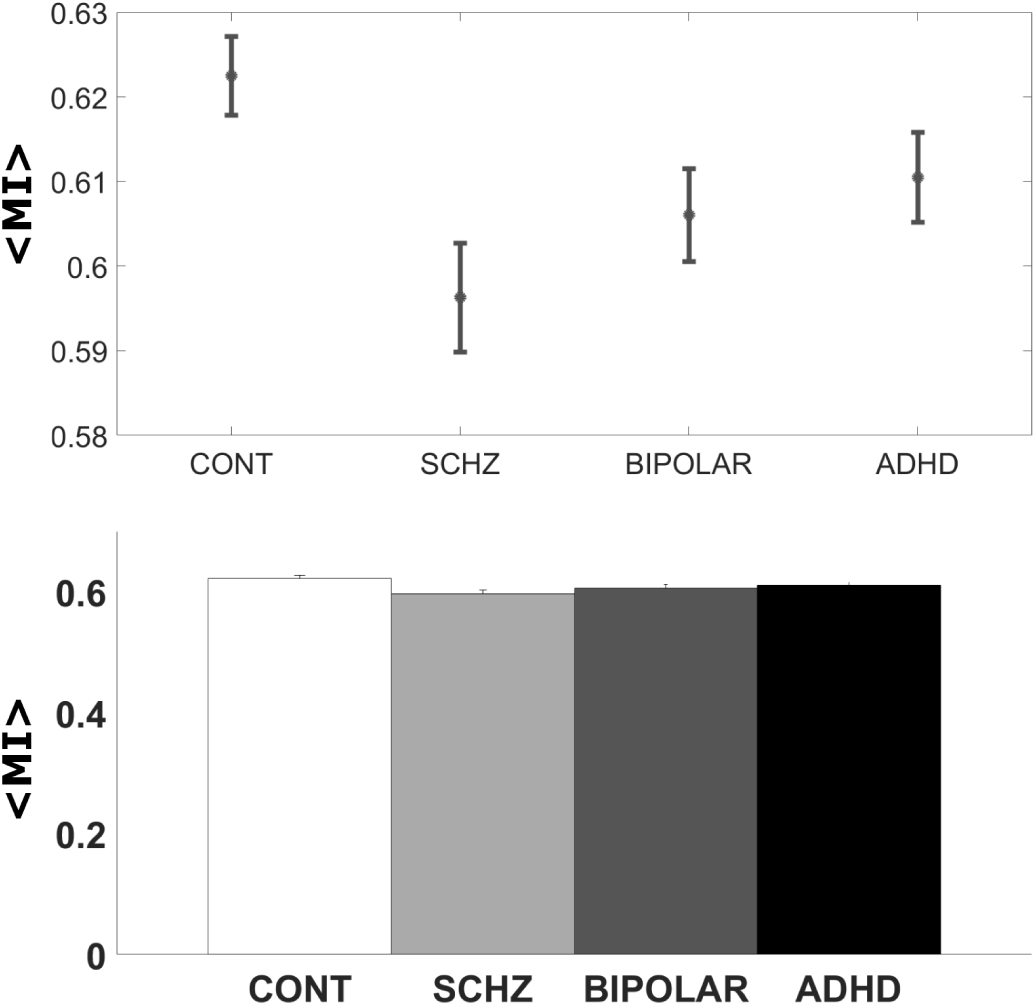
Two visually attractive but statistically suboptimal ways to report the group results, based on mean values and bars representing standard errors.

## Footnotes

1 https://github.com/danielemarinazzo/VisibilityLA5Cdata

2 https://github.com/danielemarinazzo/Visibility

3 http://www.maths.qmul.ac.uk/∼lacasa/Software.html

4 https://github.com/GRousselet/matlabstats

5 https://garstats.wordpress.com/2016/07/12/shift-function/

6 https://github.com/CPernet/RobustStatisticalToolbox/

7 https://github.com/GRousselet/matlabvisualisation

## Acknowledgments

We thank Matteo Fraschini (University of Cagliari) for setting up the Erasmus mobility for Speranza. We thank Caroline Garcia Forlim for consulting on the mutual information code. We thank Enzo Nicosia (Queen Mary University of London) for stimulating discussions on visibility graphs.

